# Enhancement of Temporal Processing via Transcutaneous Vagus Nerve Stimulation

**DOI:** 10.1101/2024.02.12.579950

**Authors:** Mehrdad Bahadori, Neha Bhutani, Simone Dalla Bella

**Author notes:** Corresponding authors, BRAMS Laboratory, Department of Psychology, University of Montreal, Montreal, Canada. Revai Inc., Montreal, Canada.

## Abstract

**Background:** The vagus nerve, a crucial component of the parasympathetic nervous system, serves as a vital communication link between the brain and body. Recent studies indicate that auricular stimulation of the vagus nerve can influence executive functions by increasing activity in brain regions like the prefrontal cortex. While prefrontal areas are associated with temporal processing, it remains unclear whether vagus nerve stimulation can also impact time perception.

**Hypothesis:** The stimulation of the vagus nerve via its auricular branch may enhance performance in temporal processing by boosting activities in prefrontal brain areas related to temporal processing.

**Methods:** Temporal processing abilities were assessed using an anisochrony detection task, where participants identified temporal irregularities in otherwise isochronous sequences while undergoing transcutaneous Vagus Nerve Stimulation (tVNS) or sham stimulation.

**Results:** The results of this study, for the first time, revealed that participants could recognize smaller temporal shifts when the vagus nerve was stimulated, compared to the sham condition.

**Conclusion:** The findings suggest that vagus nerve stimulation modulates temporal processing, supporting the notion that transcutaneous stimulation of the vagus nerve can influence cognitive functions related to temporal processing, possibly by enhancing prefrontal activities.

## Introduction

The autonomic nervous system regulates the functions of numerous organs, glands, and involuntary muscles in the body through its sympathetic and parasympathetic divisions. The Vagus nerve, the 10th and longest of the cranial nerves, is a significant component of the parasympathetic nervous system, playing a crucial role in the communication between the body and the brain. [1]. The vagus nerve extends from the brain stem to the proximal two-thirds of the colon [2] and innervates various thoracic and abdominal viscera. About 20% of vagal nerve fibers are efferent, responsible for transmitting information from the body to the brain, while the majority of remaining vagal nerve fibers are afferent fibers, sending signals from the brain to the body. [3].

In the early 1880s, American neurologist James Leonard Corning proposed invasively stimulating the vagus nerve (iVNS) using a carotid fork^1^ to treat epilepsy [4]. Due to severe side effects, this method was abandoned [5], but a century later it resurfaced as studies on animals such as cats and dogs showed that iVNS can induce synchronization or desynchronization of the electroencephalographic (EEG) signals [6], prompting more research on its impact on pharmacoresistant epilepsy [7]. Later, reported positive mood changes in patients after iVNS treatment led to clinical trials for pharmacoresistant depression [8]. In particular, the proposal of transcutaneously (non-invasively) stimulating the vagus nerve represented in the external auditory canal by Ventureyra (2000) boosted the vagal nerve stimulation studies [4, 9]. Notably, acupuncture-based stimulation in this region has demonstrated seizure control benefits [9-11].

fMRI studies on awake humans showed that transcutaneous vagus nerve stimulation (tVNS) over the auricular branch of the vagus nerve induces activity within the medulla region of the brainstem including the nucleus tractus solitarius (NTS) and to its projections such as the locus coeruleus (LC), cerebellum, amygdala, insula, nucleus accumbens, paracentral lobule of the cortex, and somatosensory cortex [12-15]. Notably, the NTS projects to the LC and dorsal raphe nucleus among others, leading to increased circulation of monoaminergic neurotransmitters (norepinephrine, serotonin, and dopamine)[1, 16, 17] as well as GABA [18, 19]. Due to these modulations, cortical areas such as the cingulate and prefrontal cortices, which are crucial for executive functions [20-23], have also been reported to exhibit increased activity [24-26]. Additionally, an increment in GABA circulation, secondary to norepinephrine surges, may promote the parasympathetic response, reducing physiological indices of stress, such as heart rate and blood pressure, which are associated with cognitive functions [27]. It is worth noting that deactivations in the hippocampus and the hypothalamus are also reported after transcutaneously stimulating the vagus nerve [13, 26].

Recent research has aimed to investigate the impact of tVNS on executive functions, given that the stimulation tends to enhance activation in brain areas such as the prefrontal cortex through the locus coeruleus-norepinephrine (LC-NE) system [28-30]. Notably, prefrontal areas such as the dorsolateral prefrontal cortex (dlPFC) play a key role in various executive functions such as cognitive flexibility, planning, inhibition, and abstract reasoning [31]. Executive functions modulated by tVNS include response inhibition [32, 33] and conflict processing [34]. For example, Sellaro et al. (2015) found that tVNS was associated with increased post-error slowing, implying an increase in response inhibition. Fischer et al. (2018) found that tVNS increased the Simon effect, measured by response accuracy and time during trials with conflict compared to non-conflict trials, thus indicating increased efficiency in conflict processing.

As tVNS increases the activity of brain areas such as prefrontal cortices, one might anticipate modifications in other cognitive abilities linked to prefrontal cortices. Other abilities that are good candidates to be affected by tVNS include the perception of event timing. [35]. Indeed, the prefrontal cortex, particularly the dlPFC, is engaged in the temporal organization of behavior (e.g. action planning) [36], mediating supra-second time estimation [37, 38], millisecond-based judgments [39-41], and other time-dependent abilities such as rhythm perception [42, 43]. Notably, the dlPFC is also activated in motor timing tasks, such as sensorimotor synchronization in finger tapping [44-46].

Despite tVNS’s capability to modulate the activity of prefrontal areas [47] involved in temporal processing, it yet remains unknown whether tVNS can also modulate performance in time perception. This study aims to test the effect of tVNS on temporal processing. To achieve this, temporal processing abilities were assessed through an anisochrony detection task, where participants were required to identify temporal irregularities in otherwise isochronous sequences while undergoing tVNS or sham stimulation. In the tVNS condition, the vagus nerve was stimulated through the auricular branch of the vagus nerve, while in the sham condition, the vagus nerve was not stimulated. Considering the role of prefrontal areas in temporal processing, it was anticipated that tVNS would enhance the detection of temporal irregularities.

## Methods

### Participants

Twenty-four young adults (17 female; 23 ± 3.7 years old) took part in the experiment. Inclusion criteria involved the absence of neurological or mental disorders, no history of brain surgery, no chronic or acute medication, non-pregnancy, no heart-related diseases, absence of metal implants in the face or brain, no pacemaker, and no piercing, implants, or physical alterations in the ear. The experimental protocol adhered to The Code of Ethics of the World Medical Association (Declaration of Helsinki) for research involving humans and received approval from the Ethics Committee for Research in Education and Psychology at the University of Montreal (CEREP--22-117-D). Participants were provided with and signed a consent form, and they were compensated for their participation in the experiment.

### Design and procedure

Each participant underwent two testing sessions, one involving tVNS stimulation and the other with sham stimulation. To ensure a randomized, single-blinded, within-subject, and crossover design, tVNS and sham sessions occurred on separate days, with a minimum one-week gap between them to prevent potential long-lasting effects of stimulation from each session. The session structure is outlined in Fig. 1. Participants were initially requested to complete the Patient Health Questionnaire (PHQ9) and the Generalized Anxiety Disorder Scale-7 (GAD7) questionnaires. [48, 49]. Following electrode placement, the individual intensity threshold for stimulation was determined for each participant [50]. Then, the participant performed the anisochrony detection task (see description below), preceded by a training session. During the anisochrony detection task, the participant fixated their gaze on a grey cross serving as the fixation point, positioned in the middle of the screen against a very light gray background. After the completion of the anisochrony detection task, the participant filled out a tVNS adverse effects questionnaire (TAEQ)[33]. The protocol was implemented and delivered to the participants using Matlab R2018a [51]. The two sessions were counterbalanced across participants.

**Fig. 1.**
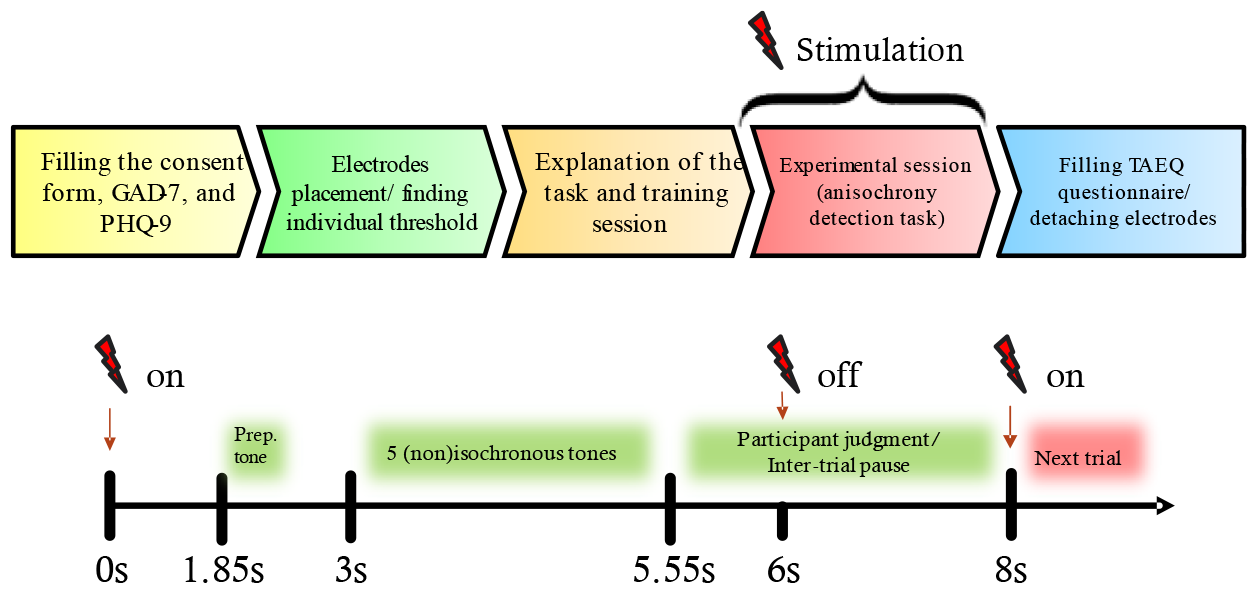
Structure of each testing session (tVNS or sham): top) steps within every session; bottom) timeline of events (in seconds) within each stimulation trial.

**Fig. 1.**
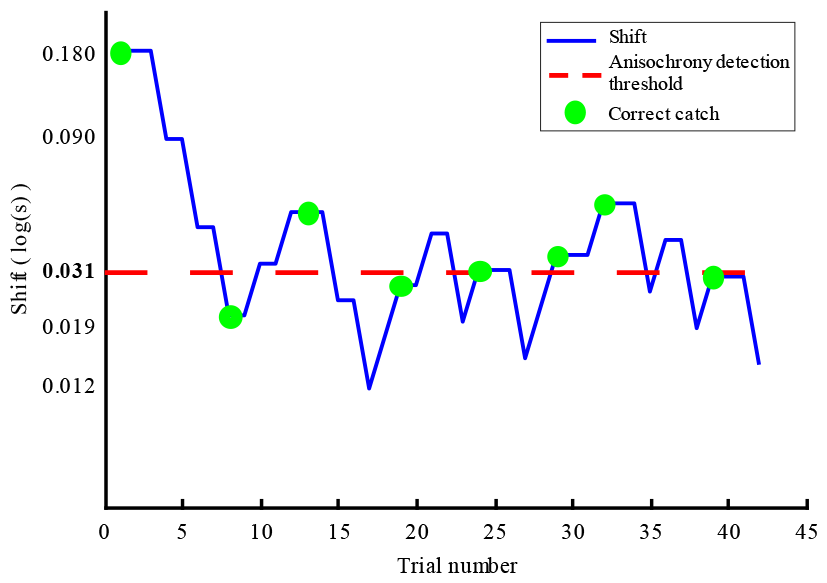
An example of a completed staircase in a single session from a participant. The y-axis represents the logarithmic scale of time. The blue line depicts temporal shifts throughout a session (reversals take place where the direction of the blue line changes). The red dashed line indicates the calculated anisochrony detection threshold, and the green circles show correct catch trials.

### Anisochrony detection task

To quantify temporal processing modulations, we adapted the anisochrony detection with tones task from the Battery for the Assessment of Auditory Sensorimotor and Timing Abilities (BAASTA)[52]. In this task, the ability to perceive a temporal irregularity (i.e., a time shift) in an isochronous sequence of tones (i.e., a metronome) was assessed. Sequences of 5 tones (tone frequency = 1047 Hz; tone duration = 150 ms) were presented. Isochronous sequences had a 600 ms constant inter-onset interval (IOI), while in non-isochronous sequences the 4th tone was delayed (shifted in time toward the 5th tone). In every trial, the participant was asked to report whether each sequence was “regular” (5 tones with the same IOI) or “irregular” (4^th^ tone was delayed), by clicking the left or right button of a computer mouse respectively. The magnitude of the temporal shift was starting at 180 ms, and the temporal shift was later controlled by a 2 correct - downward / 1 wrong - upward staircase procedure (for an example of the staircase, see Fig. 2; for an overview, refer to experiment 2 in [52]). Note that “correct” refers to detecting a non-isochronous sequence as “irregular,” and “wrong” refers to detecting a non-isochronous trial as “regular.” An up reversal was detected when the temporal shift decreased after one or more consecutive increments, and a down reversal was detected when the temporal shift increased after one or more consecutive decrements. In each trial, the metronome stimulus was preceded by a single pure tone with a lower frequency (500 Hz, 150 ms) as an alert for the start of the trial. During the staircase, 1 isochronous “catch” trial was randomly included in each set of 5 trials (1 isochronous for every 4 non-isochronous trials), for which the expected response was “regular”. These catch trials were not considered for magnitude change modifications in the staircase, but rather for stopping the staircase early if the participant frequently reported them as “irregular.” The protocol was designed to stop if one of the following conditions were met: 1) if the participant completed 12 reversals (as defined by turning points in the staircase pattern (Fig. 2)), or, 2) if more than 50% of the catch trials were answered incorrectly (this criterion was evaluated starting at the 5th catch trial). Each trial had a duration of 5.55 seconds (1850ms silence + 150ms preparatory tone + 1000ms silence + 2550ms sequence of tones) and participants had 2.45 seconds to respond to each trial, which altogether resulted in a total of 8 seconds for each trial. The metronome stimuli were delivered via headphones (Sennheiser HD201) at a self-adjusted comfortable sound pressure level. Before initiating the staircase, participants were required to complete 6 practice trials with visual feedback indicating whether they correctly reported a regular or irregular trial.

**Fig. 2.**
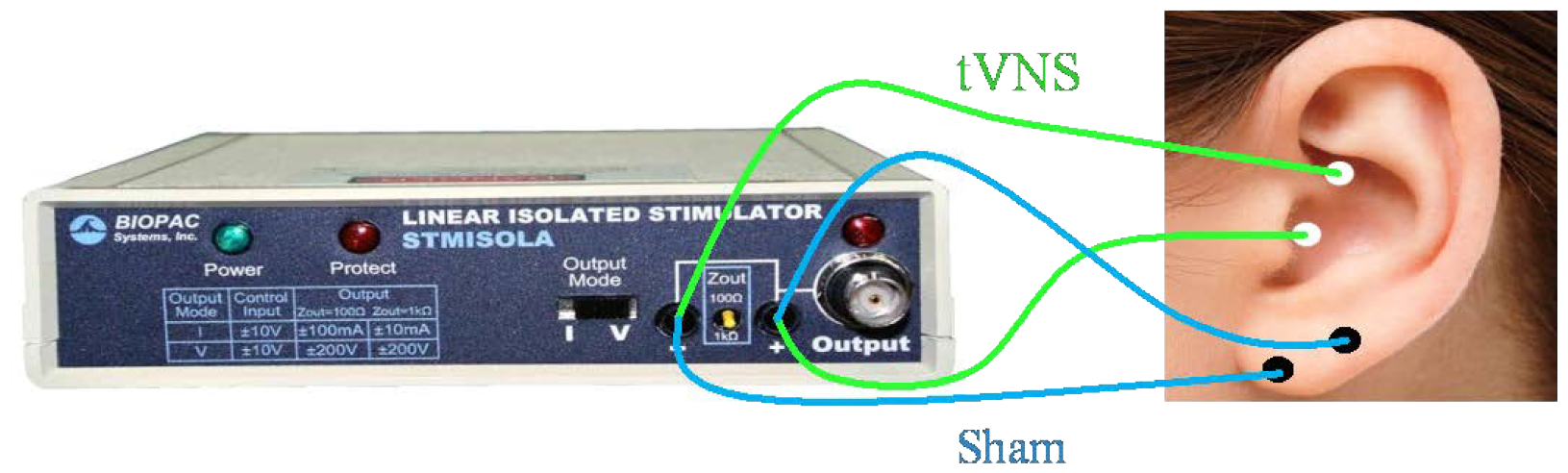
Electrode placement in tVNS and sham conditions

### tVNS

In the tVNS condition, two Ag-AgCl electrodes connected to the stimulator (STMISOLA, Biopac^®^) were attached to the left cymba conchae (Fig. 3), an area exclusively innervated by the auricular branch of the vagus nerve [53]. In the sham condition, the electrodes were placed over the left earlobe (Fig. 3), as it has been shown to be free of vagal nerves [9, 14, 53, 54]. As the thick-myelinated fibers of a sensory peripheral nerve, such as the auricular branch of the vagus nerve, mediate touch sensation, the electrical stimulus intensity for tVNS should have been adjusted to a level above the individual detection threshold and below the individual pain threshold [54]. Therefore, to determine the proper electrical stimulation intensity for each participant, two measurements were done at the minimum threshold and two at the upper level of comfort [50]. The minimum threshold was the lowest electrical intensity perceived by the participant, while the upper level of comfort was defined as the point when electrical stimulation became uncomfortable, prior to the onset of pain. The average of these four measurements was calculated and used as the electrical stimulation intensity for each participant. Also, the same process was done to determine the intensity of the stimulus for the sham condition. The stimulation was administered individually in each trial following an event-related approach [29, 55, 56], starting from the beginning of the trial and lasting for 6 seconds. The tVNS pulse width and frequency were set at 150 μs and 25Hz respectively. It is worth mentioning that side effects with these parameters are minor and mainly include skin reddening and irritation. [57, 58]. It should be mentioned that, the TAEQ included questions mesuring these possible side effects in participants [33].

**Fig. 3.**
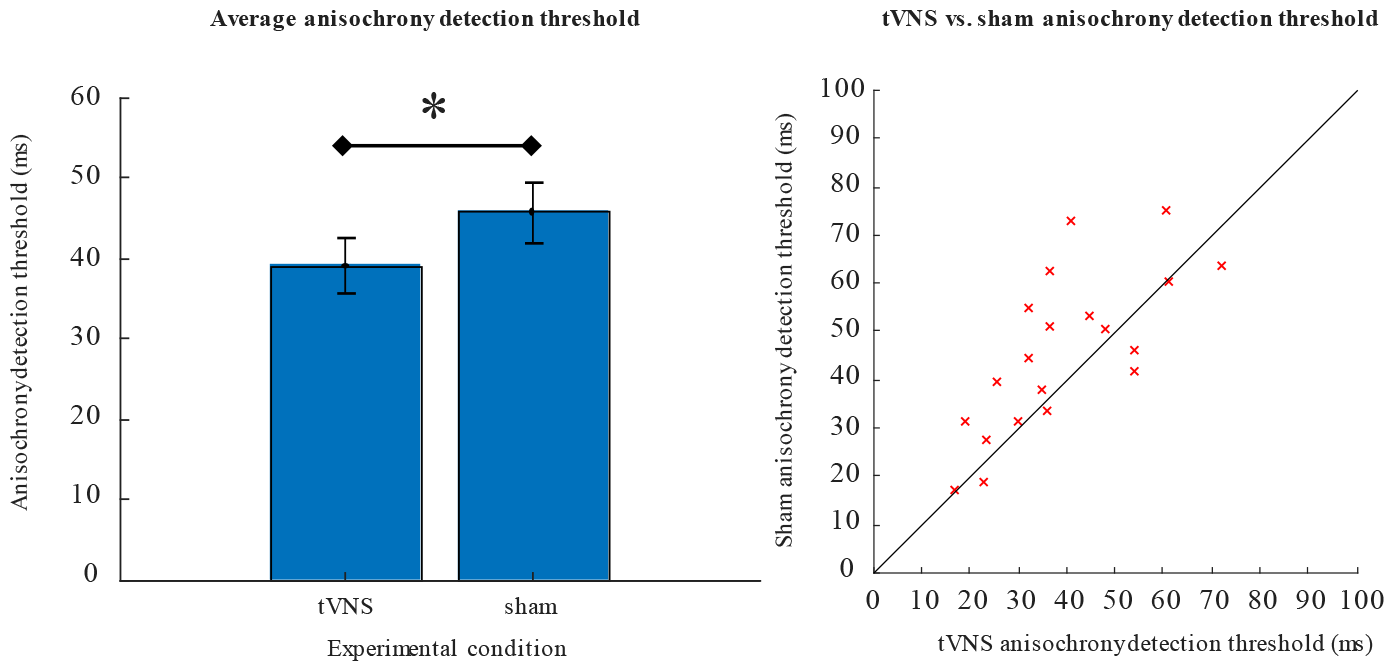
Anisochrony detection thresholds in the tVNS and the sham conditions: left panel) anisochrony detection threshold calculated by the average of the mean value of the last 6 reversals for all participants in tVNS and sham conditions. * p < 0.05; right panel) individual thresholds for tVNS vs. sham conditions in the anisochrony detection task.

### Data preparation and statistical analysis

For each session, the anisochrony detection threshold was calculated by averaging the 6 last reversal values in the staircase. IBM^®^ SPSS^®^ Statistics Version 26 [59] was then utilized to conduct a normality test, verifying the normal distribution of calculated anisochrony detection threshold values with the Shapiro-Wilk test. Therefore, thresholds between tVNS and the sham condition were compared using a paired t-test. An additional linear mixed-effects analysis was performed to study the possible effect of the duration of stimulation on detecting irregularities by time, using R Statistical Software (3.6.1.; R Core Team 2019 [60]) with the packages lmerTest [61] and lme4 [62]. In that analysis, a series of midpoint values were calculated by averaging every consecutive up and down reversal values in a staircase [63], reducing the 12 reversal values to a sequence of 6 midpoint values. Because the reversals serve as higher and lower estimates, calculating midpoints yields a more stable estimate of the response during the staircase. Then an interaction model was fit to test the effects of sequence index and stimulus condition upon midpoint values, using the lme4 formula “midpoint ∼ condition * index + (1|subject)” to specify a random intercept by subject. Sequence index values were entered as -5 to 0, i.e. centered at the final value, so that the condition coefficient would estimate the threshold at the end of the staircase.

## Results

Twenty participants (14 females; 21.7 ± 2.3 years old) were included in the final data analysis. Three participants were removed out of concern that they were answering by chance^2^. One other participant was removed because she/he was not able to reach a threshold below 70% of the initial point and visibly could not perform the task (according to the criteria presented in [64]).

We first examined whether tVNS caused a modification of the anisochrony detection threshold compared to the sham condition (see Fig. 4, left panel). It can be observed that, on average, tVNS increased detection of irregularities, indicated by lower thresholds in the tVNS condition compared to the sham condition (*t*(19) = 2.495, *p* =.022, *d* = 0.356; data were normally distributed in both conditions, as assessed with by Shapiro-Wilk test [tVNS: *p* =.485; sham: *p* = .925]). Notably, this improvement in the perception of irregularity was observed in 70% of participants (14 out of 20; Fig. 4, right panel). The average reduction in the anisochrony detection threshold for participants showing an improvement in performance due to tVNS (n = 14) was -23.3 ms ± 15.6%. To illustrate different individual responses to tVNS, Fig. 5 presents raw data and the anisochrony detection threshold from two participants who experienced a reduction in their anisochrony detection threshold under tVNS (A & B), and from two participants who did not experience any effect of tVNS (C & D), as compared to the sham condition. Accordingly, the effect of the tVNS on anisochrony detection threshold in participants A and B can be observed in the staircase trajectory as the stimulation was present throughout all the trials.

**Fig. 4.**
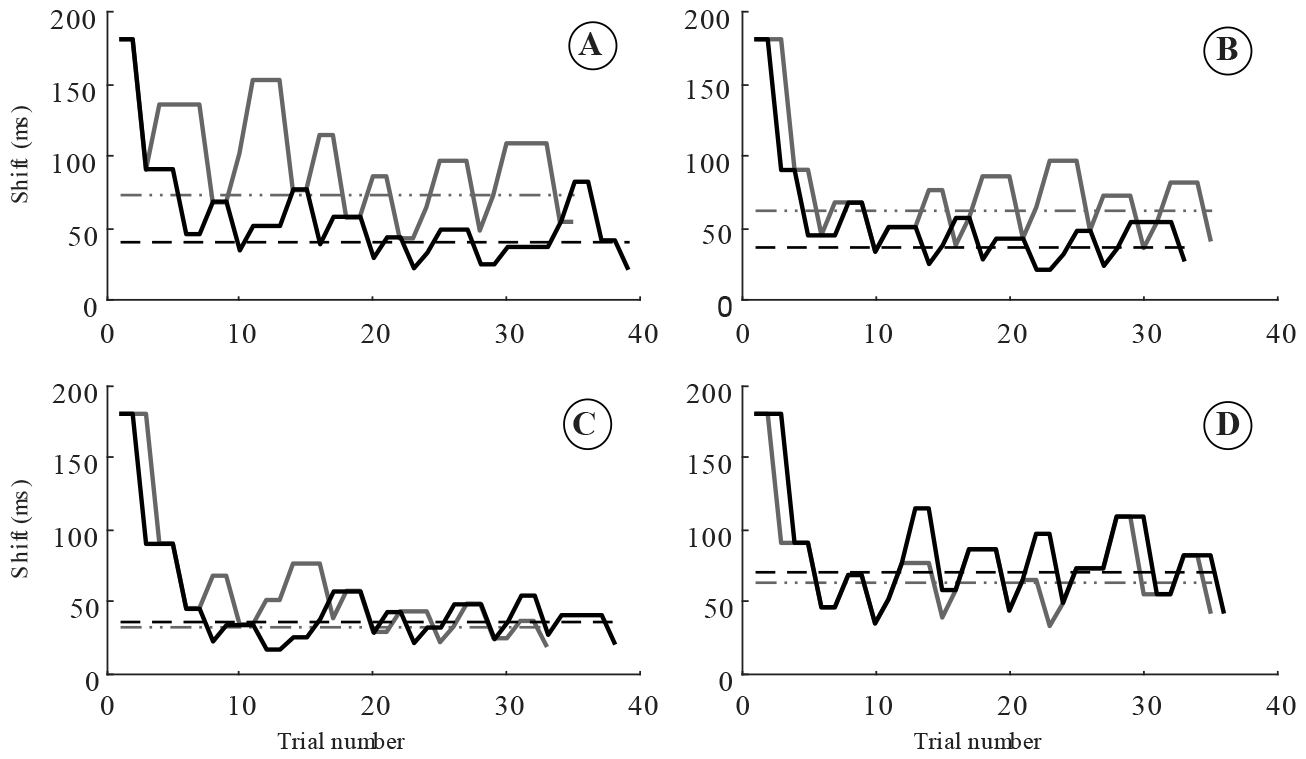
Raw staircase data and anisochrony detection thresholds for four different participants. A and B show the data for two different participants whose anisochrony detection threshold improved in the tVNS condition, while C and D show the data for two other participants whose threshold didn’t improve by tVNS. The black and grey solid lines show the temporal shifts during the anisochrony detection task in tVNS and sham conditions respectively. The black dashed line shows the anisochrony detection threshold in the tVNS condition, while the grey dash-double-dot line shows the threshold in the sham condition.

**Fig. 5.**
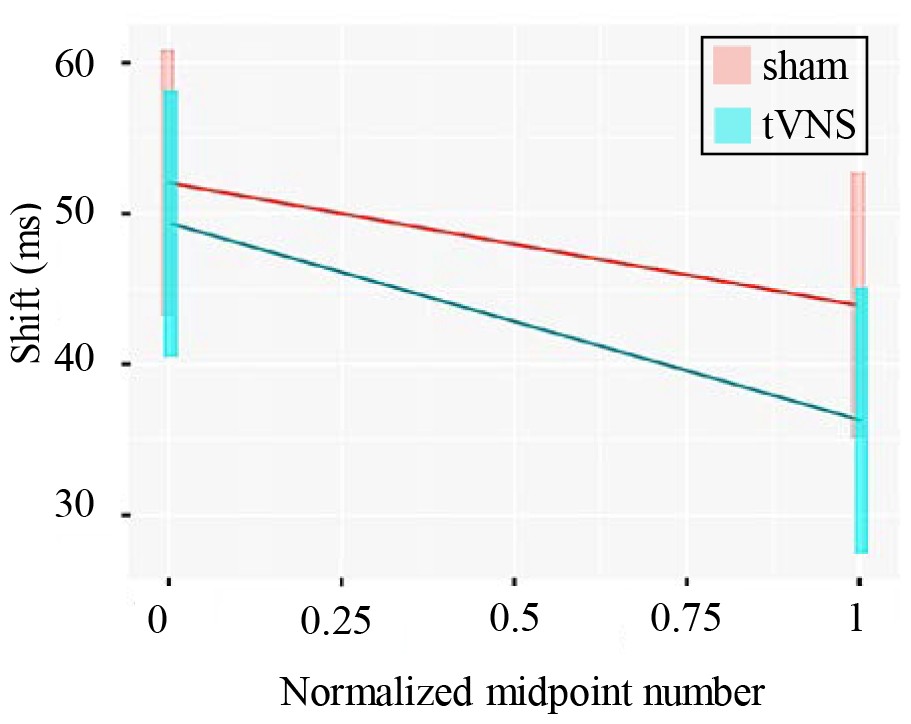
the linear prediction of the linear mixed model employed on the staircase reversal midpoint values. Note that in the x-axis, 0 corresponds to the first midpoint value, and 1 refers to the last midpoint value in the staircases.

Fig. 6 shows the linear prediction of the model for both tVNS and sham conditions. The results indicated that both the condition and the sequence index within the staircase have statistically significant effects on the staircase midpoint values. Consistent with the analysis of staircase thresholds, the omnibus tests found lower values (i.e. final reversal midpoint value in the staircase) for tVNS compared to the sham condition (F1, 217) = 10.26, p = .002). Additionally, there was a significant effect of sequence index, indicating that midpoint values decreased over the course of a staircase (F(1, 217) = 20.07, *p* < 0.001). However, this slope did not differ between conditions: there was no interaction between the effects of sequence index and condition (F(1, 217) = 1.08, *p* = .299).

We found no significant correlations between the anisochrony detection threshold and any of the GAD-7, PHQ-9, and TAEQ questionnaire data in either tVNS or sham condition (tVNS: anisochrony detection threshold vs: TAEQ: R = 0.317, p = 0.123; GAD-7: R = 0.329, p = .108; PHQ-9: R = 0.350, p = .086 – sham: anisochrony detection threshold vs: TAEQ: R = 0.366, p = 0.072; GAD-7: R = 0.327, p = .110; PHQ-9: R = 0.326, p = .112). Importantly, the GAD-7, PHQ-9, and TAEQ data in both the tVNS and sham conditions were compared to investigate potential distinctions between them. The results of the non-parametric Mann-Whitney U test (as none of both pairs (e.g., GAD-7 for tVNS and sham) were simultaneously normally distributed) revealed no significant difference between GAD-7, PHQ-9, and TAEQ data in tVNS and sham (GAD-7: *U*=197, *p*=0.947; PHQ: *U*=221, *p*=0.583; TAEQ: *U*=260, *p*=.108).

## Discussion

The present study explored the impact of tVNS on temporal processing through an anisochrony detection task, where participants were required to identify temporal shifts in an isochronous auditory stimulus. The results of both anisochrony detection threshold and linear mixed model analysis revealed for the first time that during tVNS participants were able to recognize smaller temporal shifts as compared to the sham condition. This finding suggests that tVNS modulates temporal processing, supporting the notion that the vagus nerve, when stimulated transcutaneously, can influence cognitive functions related to temporal processing.

The activation of the vagus nerve through tVNS has been demonstrated to impact diverse brain regions associated with cognitive functioning. Existing studies have documented heightened activity in cortical areas such as cingulate and prefrontal cortices, subsequent to vagus nerve stimulation [24-26]. These brain areas are known to play a crucial role in executive control, response selection, error monitoring, and conflict adaptation, which are also relevant to temporal processing [20-23].

Among the prefrontal cortices, the dorsolateral prefrontal cortex (dlPFC) has been particularly implicated in the temporal organization of behavior and cognitive processing related to time. [36]. Functional imaging studies have consistently demonstrated the involvement of dlPFC in motor timing, time estimation, and time discrimination tasks [37-41]. Moreover, dlPFC is closely associated with executive functions such as working memory, cognitive flexibility, planning, inhibition, and abstract reasoning [31]. Given the role of dlPFC in both temporal processing and executive functions, it is plausible to hypothesize that tVNS may modulate temporal perception through its effects on dlPFC and other associated brain areas.

One potential explanation for the underlying mechanisms by which tVNS influences temporal processing is the activation of the LC-NE system via the NTS, which is known to modulate cognition at various levels [28]. tVNS-induced activation of the LC-NE system may enhance the functional connectivity of brain regions involved in temporal processing, such as the dlPFC, hippocampus, and amygdala, thereby improving temporal perception [65, 66]. However, the precise mechanisms by which tVNS influences temporal perception can be further studied by using brain monitoring protocols such as fMRI or EEG.

It is worth noting that the present study focused on temporal processing in the context of anisochrony detection task, which involves the discrimination of temporal shifts. Temporal perception has been explored across various tasks, including duration discrimination, estimation, production, reproduction, temporal bisection, and detection of anisochrony, as well as the beat alignment task [67-70]. However, these different tasks are likely to involve different processes such as duration-based vs. beat-based timing, or perceptual vs. sensorimotor timing, and hence, are likely to inform us about the functioning of dissociable (or only partly overlapping) components of the timing system(s) and their associated neuronal circuitry [47]. In this regard, future research should explore the effects of tVNS on different aspects of temporal processing by deploying various tasks to gain a more comprehensive understanding of the impact of vagus nerve stimulation on time perception and temporal processing.

In the current study, the results did not show a significant correlation between the anisochrony detection threshold and the scores on the GAD-7, PHQ-9, and TAEQ questionnaires in either the tVNS or sham condition. This suggests that the observed improvement in temporal perception with tVNS was not influenced by participants’ anxiety or depression, or was caused by worsening the time performance in sham condition by putting the participant in an uncomfortable situation (e.g. inducing pain, or other feelings).

Additionally, as studies have shown activation of the LC-NE system by tVNS needs time to take effect [71, 72], one might expect to observe a significant interaction between the effects of sequence index and condition in linear mixed model analysis, which is not the case in this study. We attribute the absence of this interaction to the nature of the staircase method deployed to measure temporal processing. The staircase method inherently has a decreasing slope, starting with a relatively high value and converging downward to the desired threshold in the task [63]; thus it can potentially mask the effect of the duration of stimulation throughout reversal increments.

The application of tVNS in modulating temporal processing expands the potential uses of this non-invasive technique. Enhancing temporal processing abilities can carry significant implications across various domains. For instance, individuals with temporal processing deficits, such as those with developmental disorders like autism spectrum disorder or attention-deficit/hyperactivity disorder (ADHD), may derive benefits from interventions that improve their temporal processing skills. [73, 74]. Additionally, temporal processing plays a crucial role in tasks that require precise timing, such as music performance and speech production [75, 76], which could be potentially improved through tVNS.

## Conclusions

In conclusion, this study provides compelling evidence that tVNS effectively modulates temporal processing and time perception, as demonstrated by improved performance in an anisochrony detection task. The observed effects suggest that tVNS may enhance cognitive functions associated with temporal processing, potentially through the modulation of prefrontal cortical regions, such as dlPFC. These findings carry implications for fundamental research on temporal processing and may also inform potential therapeutic interventions for individuals with temporal processing deficits. Further investigations are warranted to explore the broader effects of tVNS on various aspects of temporal processing and to elucidate the underlying neural mechanisms.

## Submission Declaration and Verification

The authors confirm that this manuscript has not been previously published, is not under consideration elsewhere, and has the approval of all authors and responsible authorities. If accepted, it will not be published elsewhere in any form without the written consent of the copyright holder.

## Author contributions

Bahadori M: Conceptualization; Data acquisition and curation; Formal analysis; Funding acquisition; Investigation; Methodology; Project administration; Software; Supervision; Validation; Visualization; Writing - original draft - review & editing; Bhutani N: Conceptualization; Funding acquisition; Investigation; Methodology; Project administration; Resources; Supervision; Validation; Writing - review & editing; Dalla Bella S: Conceptualization; Funding acquisition; Investigation; Methodology; Project administration; Resources; Supervision; Validation; Writing - review & editing.

## Authorship

All authors should have made substantial contributions to all of the following: (1) the conception and design of the study, or acquisition of data, or analysis and interpretation of data, (2) drafting the article or revising it critically for important intellectual content, (3) final approval of the version to be submitted.

## Acknowledgment

Special thanks to Dr. Nicholas E.V. Foster for his invaluable contributions to the design of the methodology, data analysis, and the review and editing of the manuscript.

## Footnotes

1 The carotid fork was a device equipped with electrodes aimed to directly stimulate cervical branches of the vagus nerve, which are positioned next to the carotid artery in the neck.

2 These participants, in either tVNS or sham condition, exceeded the total of 50 trials, had more than 20% wrong catch trials, and have reached a minimum reversal value of smaller than 7.1 ms (2 out 3 reached below 2 ms), which was smaller than mean minus standard deviation of the minimum reversal value in all the participants in both conditions (tVNS: 10.8 ms; sham: 9.5 ms).

## References

[1] Butt MF, Albusoda A, Farmer AD, Aziz Q. The anatomical basis for transcutaneous auricular vagus nerve stimulation. Journal of anatomy 2020;236(4):588–611.

[2] Komisaruk BR, Frangos E. Vagus nerve afferent stimulation: projection into the brain, reflexive physiological, perceptual, and behavioral responses, and clinical relevance. Autonomic Neuroscience 2022;237:102908.

[3] Bonaz B, Bazin T, Pellissier S. The vagus nerve at the interface of the microbiota-gut-brain axis. Frontiers in neuroscience 2018;12:49.

[4] Colzato L, Beste C. A literature review on the neurophysiological underpinnings and cognitive effects of transcutaneous vagus nerve stimulation: challenges and future directions. Journal of neurophysiology 2020;123(5):1739–55.

[5] Lanska DJ. JL Corning and vagal nerve stimulation for seizures in the 1880s. Neurology 2002;58(3):452–9.

[6] Rutecki P. Anatomical, physiological, and theoretical basis for the antiepileptic effect of vagus nerve stimulation. Epilepsia 1990;31:S1–S6.

[7] Penry JK, Dean JC. Prevention of intractable partial seizures by intermittent vagal stimulation in humans: preliminary results. Epilepsia 1990;31:S40–S3.

[8] Cristancho P, Cristancho MA, Baltuch GH, Thase ME, John P. Effectiveness and safety of vagus nerve stimulation for severe treatment-resistant major depression in clinical practice after FDA approval: outcomes at 1 year. The Journal of clinical psychiatry 2011;72(10):5594.

[9] Ventureyra EC. Transcutaneous vagus nerve stimulation for partial onset seizure therapy. Child’s Nervous System 2000;16(2):101–2.

[10] Chen S, Wang S, Rong P, Liu J, Zhang H, Zhang J. Acupuncture for refractory epilepsy: role of thalamus. Evidence-Based Complementary and Alternative Medicine 2014;2014.

[11] Cheuk DK, Wong V. Acupuncture for epilepsy. Cochrane Database of Systematic Reviews 2014(5).

[12] Kraus T, Kiess O, Hösl K, Terekhin P, Kornhuber J, Forster C. CNS BOLD fMRI effects of sham-controlled transcutaneous electrical nerve stimulation in the left outer auditory canal–a pilot study. Brain stimulation 2013;6(5):798–804.

[13] Frangos E, Ellrich J, Komisaruk BR. Non-invasive access to the vagus nerve central projections via electrical stimulation of the external ear: fMRI evidence in humans. Brain stimulation 2015;8(3):624–36.

[14] Yakunina N, Kim SS, Nam E-C. Optimization of transcutaneous vagus nerve stimulation using functional MRI. Neuromodulation: technology at the neural interface 2017;20(3):290–300.

[15] Sclocco R, Garcia RG, Kettner NW, Isenburg K, Fisher HP, Hubbard CS, et al. The influence of respiration on brainstem and cardiovagal response to auricular vagus nerve stimulation: a multimodal ultrahigh-field (7T) fMRI study. Brain stimulation 2019;12(4):911–21.

[16] Ventura-Bort C, Wirkner J, Genheimer H, Wendt J, Hamm AO, Weymar M. Effects of transcutaneous vagus nerve stimulation (tVNS) on the P300 and alpha-amylase level: a pilot study. Frontiers in Human Neuroscience 2018;12:202.

[17] Warren CM, Tona KD, Ouwerkerk L, Van Paridon J, Poletiek F, van Steenbergen H, et al. The neuromodulatory and hormonal effects of transcutaneous vagus nerve stimulation as evidenced by salivary alpha amylase, salivary cortisol, pupil diameter, and the P3 event-related potential. Brain stimulation 2019;12(3):635–42.

[18] Capone F, Assenza G, Di Pino G, Musumeci G, Ranieri F, Florio L, et al. The effect of transcutaneous vagus nerve stimulation on cortical excitability. Journal of Neural Transmission 2015;122(5):679–85.

[19] Keute M, Ruhnau P, Heinze H-J, Zaehle T. Behavioral and electrophysiological evidence for GABAergic modulation through transcutaneous vagus nerve stimulation. Clinical Neurophysiology 2018;129(9):1789–95.

[20] Aston-Jones G, Cohen JD. An integrative theory of locus coeruleus-norepinephrine function: adaptive gain and optimal performance. Annu Rev Neurosci 2005;28:403–50.

[21] Logue SF, Gould TJ. The neural and genetic basis of executive function: attention, cognitive flexibility, and response inhibition. Pharmacology Biochemistry and Behavior 2014;123:45–54.

[22] Ullsperger M, Fischer AG, Nigbur R, Endrass T. Neural mechanisms and temporal dynamics of performance monitoring. Trends in cognitive sciences 2014;18(5):259–67.

[23] Menon V, D’Esposito M. The role of PFC networks in cognitive control and executive function. Neuropsychopharmacology 2022;47(1):90–103.

[24] Badran BW, Mithoefer OJ, Summer CE, LaBate NT, Glusman CE, Badran AW, et al. Short trains of transcutaneous auricular vagus nerve stimulation (taVNS) have parameter-specific effects on heart rate. Brain Stimulation 2018;11(4):699–708.

[25] Dietrich S, Smith J, Scherzinger C, Hofmann-Preiß K, Freitag T, Eisenkolb A, et al. A novel transcutaneous vagus nerve stimulation leads to brainstem and cerebral activations measured by functional MRI/Funktionelle Magnetresonanztomographie zeigt Aktivierungen des Hirnstamms und weiterer zerebraler Strukturen unter transkutaner Vagusnervstimulation. 2008.

[26] Frangos E, Komisaruk BR. Access to vagal projections via cutaneous electrical stimulation of the neck: fMRI evidence in healthy humans. Brain stimulation 2017;10(1):19–27.

[27] Breit S, Kupferberg A, Rogler G, Hasler G. Vagus nerve as modulator of the brain–gut axis in psychiatric and inflammatory disorders. Frontiers in psychiatry 2018:44.

[28] Poe GR, Foote S, Eschenko O, Johansen JP, Bouret S, Aston-Jones G, et al. Locus coeruleus: a new look at the blue spot. Nature Reviews Neuroscience 2020;21(11):644–59.

[29] Villani V, Finotti G, Di Lernia D, Tsakiris M, Azevedo RT. Event-related transcutaneous vagus nerve stimulation modulates behaviour and pupillary responses during an auditory oddball task. Psychoneuroendocrinology 2022;140:105719.

[30] Sargin D, Jeoung H-S, Goodfellow NM, Lambe EK. Serotonin regulation of the prefrontal cortex: cognitive relevance and the impact of developmental perturbation. ACS chemical neuroscience 2019;10(7):3078–93.

[31] Diamond A. Executive functions. Annual review of psychology 2013;64:135.

[32] Steenbergen L, Sellaro R, Stock A-K, Verkuil B, Beste C, Colzato LS. Transcutaneous vagus nerve stimulation (tVNS) enhances response selection during action cascading processes. Eur Neuropsychopharmacol 2015;25(6):773–8.

[33] Sellaro R, van Leusden Jw, Tona K-D, Verkuil B, Nieuwenhuis S, Colzato LS. Transcutaneous vagus nerve stimulation enhances post-error slowing. Journal of cognitive neuroscience 2015;27(11):2126–32.

[34] Fischer R, Ventura-Bort C, Hamm A, Weymar M. Transcutaneous vagus nerve stimulation (tVNS) enhances conflict-triggered adjustment of cognitive control. Cognitive, Affective, & Behavioral Neuroscience 2018;18:680–93.

[35] Mangels JA, Ivry RB, Shimizu N. Dissociable contributions of the prefrontal and neocerebellar cortex to time perception. Cognitive Brain Research 1998;7(1):15–39.

[36] Fuster JM. Temporal processing. Annals of the New York Academy of Sciences 1995;769:173–81.

[37] Basso G, Nichelli P, Wharton CM, Peterson M, Grafman J. Distributed neural systems for temporal production: a functional MRI study. Brain research bulletin 2003;59(5):405–11.

[38] Macar F, Lejeune H, Bonnet M, Ferrara A, Pouthas V, Vidal F, et al. Activation of the supplementary motor area and of attentional networks during temporal processing. Experimental Brain Research 2002;142(4):475–85.

[39] Rao SM, Harrington DL, Haaland KY, Bobholz JA, Cox RW, Binder JR. Distributed neural systems underlying the timing of movements. Journal of Neuroscience 1997;17(14):5528–35.

[40] Maquet P, Lejeune H, Pouthas V, Bonnet M, Casini L, Macar F, et al. Brain activation induced by estimation of duration: a PET study. Neuroimage 1996;3(2):119–26.

[41] Smith A, Taylor E, Lidzba K, Rubia K. A right hemispheric frontocerebellar network for time discrimination of several hundreds of milliseconds. NeuroImage 2003;20(1):344–50.

[42] Thaut MH, Trimarchi PD, Parsons LM. Human brain basis of musical rhythm perception: common and distinct neural substrates for meter, tempo, and pattern. Brain sciences 2014;4(2):428–52.

[43] Thaut MH. Neural basis of rhythmic timing networks in the human brain. Annals of the New York Academy of Sciences 2003;999(1):364–73.

[44] Larsson J, Gulyás B, Roland PE. Cortical representation of self-paced finger movement. Neuroreport 1996;7(2):463–8.

[45] Rubia K, Overmeyer S, Taylor E, Brammer M, Williams S, Simmons A, et al. Prefrontal involvement in temporal bridging and timing movement. Neuropsychologia 1998;36(12):1283–93.

[46] Rubia K, Overmeyer S, Taylor E, Brammer M, Williams S, Simmons A, et al. Functional frontalisation with age: mapping neurodevelopmental trajectories with fMRI. Neuroscience & Biobehavioral Reviews 2000;24(1):13–9.

[47] Rubia K, Smith A. The neural correlates of cognitive time management: a review. Acta neurobiologiae experimentalis 2004.

[48] Kroenke K, Spitzer RL, Williams JB. The PHQ-9: validity of a brief depression severity measure. Journal of general internal medicine 2001;16(9):606–13.

[49] Spitzer RL, Kroenke K, Williams JB, Löwe B. A brief measure for assessing generalized anxiety disorder: the GAD-7. Archives of internal medicine 2006;166(10):1092–7.

[50] Thakkar VJ, Engelhart AS, Khodaparast N, Abadzi H, Centanni TM. Transcutaneous auricular vagus nerve stimulation enhances learning of novel letter-sound relationships in adults. Brain Stimulation 2020;13(6):1813–20.

[51] Inc. TM. MATLAB version: 9.4.0 (R2018a). The MathWorks Inc.; 2018.

[52] Dalla Bella S, Farrugia N, Benoit C-E, Begel V, Verga L, Harding E, et al. BAASTA: battery for the assessment of auditory sensorimotor and timing abilities. Behavior Research Methods 2017;49(3):1128–45.

[53] Peuker ET, Filler TJ. The nerve supply of the human auricle. Clinical Anatomy 2002;15(1):35–7.

[54] Ellrich J. Transcutaneous auricular vagus nerve stimulation. Journal of Clinical Neurophysiology 2019;36(6):437–42.

[55] Sharon O, Fahoum F, Nir Y. Transcutaneous vagus nerve stimulation in humans induces pupil dilation and attenuates alpha oscillations. Journal of Neuroscience 2021;41(2):320–30.

[56] Llanos F, McHaney JR, Schuerman WL, Han GY, Leonard MK, Chandrasekaran B. Non-invasive peripheral nerve stimulation selectively enhances speech category learning in adults. NPJ science of learning 2020;5(1):1–11.

[57] Hilz MJ. Transcutaneous vagus nerve stimulation-A brief introduction and overview. Autonomic Neuroscience 2022:103038.

[58] Badran BW, Alfred BY, Adair D, Mappin G, DeVries WH, Jenkins DD, et al. Laboratory administration of transcutaneous auricular vagus nerve stimulation (taVNS): technique, targeting, and considerations. JoVE (Journal of Visualized Experiments) 2019(143):e58984.

[59] IBM Corp. Released 2019. IBM SPSS Statistics for Windows VA, NY: IBM Corp.

[60] 3.6.1.; R Core Team 2019.

[61] Kuznetsova A, Brockhoff PB, Christensen RHB. lmerTest package: tests in linear mixed effects models. Journal of statistical software 2017;82(13).

[62] Bates D, Kliegl R, Vasishth S, Baayen H. Parsimonious mixed models. arXiv preprint arXiv:150604967 2015.

[63] Levitt H. Transformed up-down methods in psychoacoustics. The Journal of the Acoustical society of America 1971;49(2B):467–77.

[64] Oschkinat M, Hoole P, Falk S, Dalla Bella S. Temporal malleability to auditory feedback perturbation is modulated by rhythmic abilities and auditory acuity. Frontiers in Human Neuroscience 2022;16.

[65] Bahtiyar S, Karaca KG, Henckens MJ, Roozendaal B. Norepinephrine and glucocorticoid effects on the brain mechanisms underlying memory accuracy and generalization. Molecular and Cellular Neuroscience 2020;108:103537.

[66] Broncel A, Bocian R, Kłos-Wojtczak P, Kulbat-Warycha K, Konopacki J. Vagal nerve stimulation as a promising tool in the improvement of cognitive disorders. Brain research bulletin 2020;155:37–47.

[67] Dalla Bella S, Sowiński J. Uncovering beat deafness: detecting rhythm disorders with synchronized finger tapping and perceptual timing tasks. JoVE (Journal of Visualized Experiments) 2015(97):e51761.

[68] Merchant H, De Lafuente V. Introduction to the neurobiology of interval timing. Springer; 2014.

[69] Sowiński J, Dalla Bella S. Poor synchronization to the beat may result from deficient auditory-motor mapping. Neuropsychologia 2013;51(10):1952–63.

[70] Grondin S. Timing and time perception: A review of recent behavioral and neuroscience findings and theoretical directions. Attention, Perception, & Psychophysics 2010;72(3):561–82.

[71] Neuser MP, Teckentrup V, Kühnel A, Hallschmid M, Walter M, Kroemer NB. Vagus nerve stimulation boosts the drive to work for rewards. Nature communications 2020;11(1):1–11.

[72] Sun J-B, Cheng C, Tian Q-Q, Yuan H, Yang X-J, Deng H, et al. Transcutaneous Auricular Vagus Nerve Stimulation Improves Spatial Working Memory in Healthy Young Adults. Frontiers in neuroscience 2021;15.

[73] Allman MJ. Deficits in temporal processing associated with autistic disorder. Frontiers in integrative neuroscience 2011;5:2.

[74] Hart H, Radua J, Mataix-Cols D, Rubia K. Meta-analysis of fMRI studies of timing in attention-deficit hyperactivity disorder (ADHD). Neuroscience & Biobehavioral Reviews 2012;36(10):2248–56.

[75] Wittmann M, Pöppel E. Temporal mechanisms of the brain as fundamentals of communication—with special reference to music perception and performance. Musicae Scientiae 1999;3(1_suppl):13–28.

[76] Brown S, Pfordresher PQ, Chow I. A musical model of speech rhythm. Psychomusicology: Music, Mind, and Brain 2017;27(2):95.

